# Liquid-solid coexistence at single fibril resolution during tau condensate ageing

**DOI:** 10.1101/2025.02.06.636870

**Authors:** Mahaprasad S. R. Sahu, Jiapeng Wei, David A. Weitz, Tuomas P. J. Knowles, Kanchan Garai

## Abstract

Aggregated fibrillar tangles of the microtubule-associated protein tau are a hallmark of Alzheimer’s disease. It is becoming increasingly clear that a key process that can trigger the formation of such tau tangles is the aberrant ageing of biomolecular condensates of tau formed via liquid-liquid phase separation. This ageing process affects the overall mechanical and structural properties of the condensates, but the molecular-level mechanisms by which aggregation takes place in the condensates have remained elusive. Here, by tracking individual tau molecules inside the condensates using single molecule microscopy, we show that the ageing process is characterized by the coexistence of a growing solid phase of tau within a dense liquid phase. The liquid phase is increasingly confined to the pores of the growing fibril gel network, but maintains its initial viscosity. These findings add a spatial dimension to the ageing of condensates and demonstrate that spatial heterogeneity is a key feature of the liquid-to-solid transition of tau.

## Introduction

The microtubule-associated protein tau is involved in the pathology of multiple neurodegenerative disorders, commonly classified as tauopathies. The spectrum of tauopathies includes Alzheimer’s disease, Pick’s disease and progressive supranuclear palsy etc.^1–3^ A pathological hallmark in all of these diseases is the accumulation of intracellular filaments of tau in the neuronal cells in the brain. These filaments, commonly known as neurofibrillary tangles bear characteristics of the amyloid fibrils. However, the individual micro-steps involved in the reaction chain starting from soluble monomers to the insoluble fibrils are not well understood. The canonical pathway of aggregation of tau involves primary nucleation, followed by elongation and self-replication of the amyloid fibrils (Figure 1A).^4,5^ The rate constants of these processes have been estimated from the kinetic studies through the application of the integrated rate laws. ^6,7^ Recent studies, however, indicate that like many other intrinsically disordered proteins (IDPs), tau undergoes liquid-liquid phase separation (LLPS) both in vitro and in vivo.^8^ The functional role of the tau condensates is increasingly linked to control of the cytoskeleton where the liquid tau droplets facilitate nucleation and stabilization of the microtubules in the axons of the neuronal cells.^9,10^ While LLPS of proteins and protein-nucleic acid complexes are emerging as a fundamental principle underpinning the formation of membrane-less organelles performing multitude of cellular functions, ^11^ it has a dark side too. Protein-rich condensates arising from LLPS are metastable and, hence undergo irreversible liquid-to-solid transition (LST) in the form of hydrogels^12^ and amyloid fibrils. Mounting evidence suggests that the aberrant liquid-solid transition of the condensates, for instance of *α*-synuclein,^13^ TDP-43,^14^ FUS,^15^ and tau,^16,17^ is linked to the pathogenesis of various neurodegenerative diseases.^18^ This non-canonical mechanism of protein aggregation where fibrillization occurs via LLPS remains poorly understood. For example, the rates of nucleation or growth of the fibrils formed in this pathway have remained challenging to quantify.

**Figure 1:**
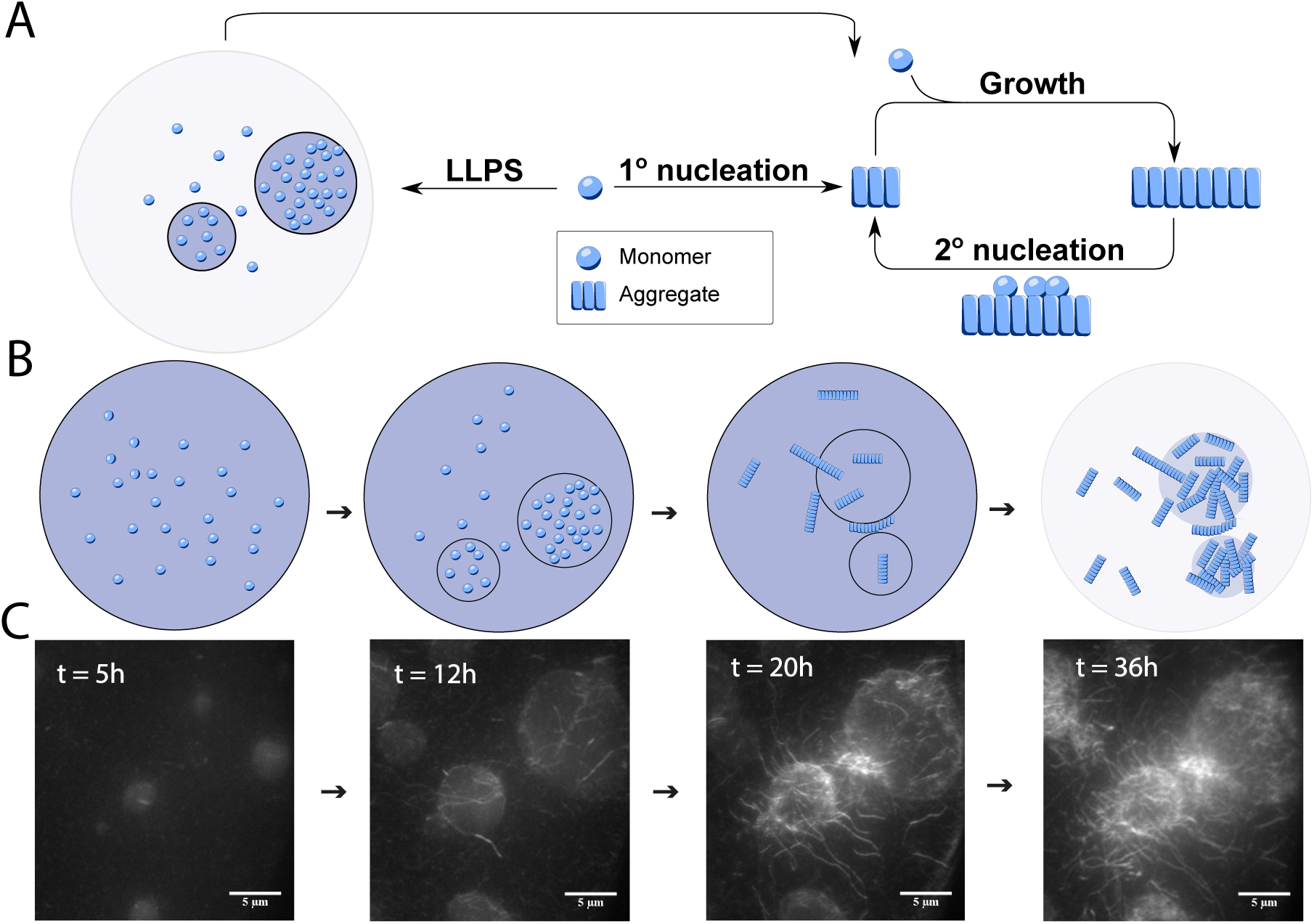
Aggregation reaction network and phase transitions of tau. (A) Tau LLPS and aggregation reaction network, comprising: primary nucleation, elongation and secondary nucleation. (B) Liquid-liquid phase separation and liquid-to solid transition of P301L tau + polyU RNA. Fibrils can form inside, outside, or partly inside and partly outside of the condensates. (C) TIRF images of 10 *µM* P301L tau in the presence of 40 *µg/ml* polyU at *t* = 5, 12, 20, 36 hours, exhibiting the three different stages outlined in B: Homogeneous solution phase, LLPS and LLPS to fibril transition.

Studies using full-length and various truncated constructs of tau have revealed that the LLPS of tau is primarily governed by weak but multivalent interactions.^19–21^ The liquid-to-solid transition, by contrast, leads to amyloid-like structures which are stabilized by backbone-backbone hydrogen bonding interactions. Specific point mutations such as P301L, P301S, G272V, and ΔK280^22^ that are involved in various tauopathies accelerate the liquid-solid transition considerably. Here, we use fluorescence microscopy to visualize the liquid condensates and the amyloid fibrils of tau and attempt to quantify the rates of the crucial reaction steps such as primary nucleation and elongation of the fibrils.

Primary nucleation is the first step in the kinetics of fibrillization. However, it is an intrinsically slow event at physiological concentrations of the protein due to the high kinetic barrier arising from the high cost of surface energy required to create a nucleus.^23–25^ Therefore, experimental measurement of primary nucleation is lacking. However, in the event of LLPS, the concentration of the protein in the protein-dense phase can be very high. Hence, it can facilitate primary nucleation by lowering the kinetic barrier.^26^ The measurement of primary nucleation initiating the liquid-to-solid transition of the condensates, however, has not yet been done due to the lack of suitable techniques. We posit that single-molecule fluorescence techniques offer the opportunity to study nucleation and replication of the fibrils at the single aggregate level. ^27,28^

Here, we use total internal reflection fluorescence (TIRF) microscopy and spinning disk super resolution by optical pixel reassignment (SoRa) microscopy to monitor the time evolution of the condensates and the growth of single fibrils of tau within and outside the condensates. We find that the condensates, particularly the interface of the condensates, facilitate nucleation of the amyloid fibrils. These fibrils coexist and expand within and outside of the liquid phase. We monitored the viscosity and surface tension of the liquid phase of the condensates using single-particle tracking, fluorescence recovery after photobleaching (FRAP), and fusion kinetic measurements to examine spatial and temporal changes in condensate structure and dynamics. The viscosity and surface tension of the dense liquid phase remain constant with ageing time. Upon ageing both the liquid and the solid phases coexist within the same condensate, leading to structures that are highly heterogeneous.^29^ Although the volume fraction of the liquid phase diminishes during aging, the viscosity remains nearly unchanged.

## Results and discussion

We focus on the P301L variant of tau to study the phase transitions from the homogeneously mixed state to the LLPS state and subsequently to the solid amyloid state. P301L tau was chosen since it undergoes liquid-to-solid transition within a few hours, unlike the WT-tau, which may take several days. Importantly, the mutation P301L is also associated with a higher risk of frontotemporal dementia.^10,28^ To induce LLPS we add 40 *µg/ml* polyU RNA to the solutions of 5 to 40 *µM* tau. The appearance of the liquid condensates and the fibrillar aggregates is monitored by TIRF microscopy with ThT as the reporting dye. Therefore, these experiments are performed using unlabeled tau without any covalent modification of the protein.

### P301L tau with polyU undergoes liquid liquid phase separation and liquid-to-solid transition in vitro

Earlier studies have shown that, above a critical concentration, a homogeneous tau solution undergoes phase separation, leading to the formation of protein-dense liquid droplets that coexist with the dilute phase.^16,19^ The liquid droplets undergo irreversible conversion to amyloid fibrils upon aging. The LLPS and the liquid-to-solid processes are schematically shown in Figure 1B. The TIRF microscopy images recorded at various time points demonstrate these three stages, as depicted in Figure 1C. Initially, at *t* = 0 − 4 *h*, the system is homogeneous. After 5 hours, liquid droplets appear and continue to grow via fusion and Oswald ripening,^20,30^ resulting in the formation of bright and round condensates. At *t* = 12 − 20 *h*, the formation of fibrils is evident both within and outside of the condensates, with some fibrils appearing partially within the condensate and partially outside of it. By *t* = 36 *h*, the condensates fully solidify into fibrillar aggregates, which protrude into the dilute phase, therefore, the boundary between the dense and dilute phases becomes more indistinct. Akin to the aggregation of most of the amyloid proteins, aggregation of tau involves the following microscopic steps: primary nucleation, elongation, and secondary processes.^27^ Figure 1A shows the reaction network of tau LLPS and tau aggregation. Since secondary process events are difficult to quantify,^31^ here we measured the rates of the primary nucleation events and the elongation of the fibrils of P301L tau.

### Direct observation of the elongation of P301L tau fibrils

To study LLPS and aggregation kinetics at the single-aggregate level, we captured TIRF microscopy images at intervals of 30 minutes at the same position of the sample for about 24 hours. The videos show clear evidence of elongation of the fibrils both in the dilute and the dense phases. The red lines, automatically traced fibrils, are superimposed on a TIRF microscopy image sequence captured at various time points, as shown in Figure 2A-C. Careful examination of the videos reveals a distinct growth pattern for the fibrils: initially, the fibrils exhibit linear growth, followed by a pause, and subsequently resumption of the growth at a similar rate (Supplementary Video 1). This stop-and-go pattern of the fibrillar growth has been documented previously in the case of *Aβ*40 and *Aβ*42 in the absence of LLPS.^31,32^ The rates of elongation are estimated from the plots of the length of the fibrils versus time after removing the paused sections. Figure 2D shows the length versus time plot of the fibrils (n = 69) tracked in the dilute phase in the case of 10 *µM* P301L tau. The distribution of the rates of elongation resembles a log-normal distribution (Figure 2E) with a peak at 4 *µm/h*. A similar distribution for the rate of elongation has been reported earlier in the case of *Aβ*40 and *Aβ*42.^31–33^ Similarly, the fibrils inside the condensates were also traced and analyzed (Supplementary Figure S1). We find that the rate of elongation of the fibrils is nearly the same within the range of concentration of tau used here, i.e., for 5, 7.5, 10 and 15 *µM* (Figure 2F and Supplementary Figure S1). This may be expected since concentrations of the protein in the dilute and the dense phases are expected to remain invariant upon phase separation, although the volume fraction of the dense phase increases with the increase of the initial concentration. Surprisingly, the rate of elongation of the fibrils in the dense phase is found to be same as that in the dilute phase, even though the concentrations of the soluble tau differ by 2 orders of magnitude between the two phases (Supplementary Figure S2). The rate constant of a reaction in a crowded environment in the presence of a high concentration of reactant molecules depends on multiple factors, including excluded volume effects, hydrodynamic interactions, diffusivity of the reactant molecules and caging effects.^34,35^ Later we will measure the diffusivity of the tau molecules inside the condensates to account for its role in affecting the growth of the fibrils.

**Figure 2:**
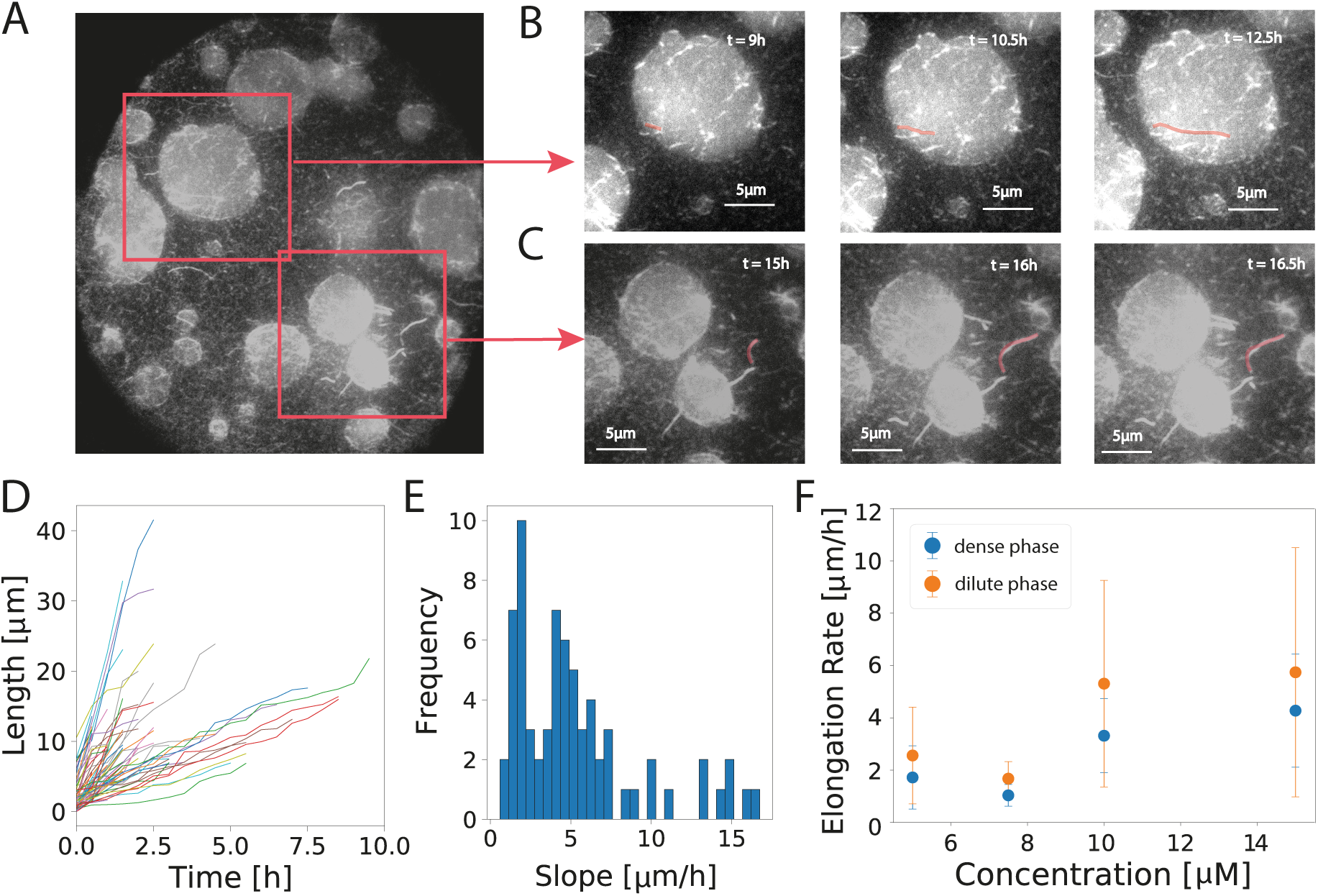
Elongation of the fibrils in the dilute and the dense phases. (A) TIRF image recorded with 10*µM* of P301L tau in presence of 40 *µg/ml* polyU RNA. (B) Tracked fibrils in the dense phase (red) overlaid on a TIRFM image sequence at 9, 10.5, 12.5 *h*. (C) Tracked fibrils in the dilute phase (red) overlaid on a TIRFM image sequence at 15, 16, 16.5 *h*. (D) Time traces of the individual fibrils. The paused sections, due to the stop-and-go elongation behavior, have been removed. (E) Histogram of the elongation rates estimated from D. (F) Average elongation rates measured at 5 *µM* to 15 *µM* of P301L tau + 40 *µg/ml* polyU RNA. Error bars represent ± standard deviation.

### Direct observation of nucleation in condensates

Then we investigate whether and how the liquid condensates of tau can facilitate nucleation of the fibrils. Indeed we observe numerous events of nucleation inside the condensates in the solution of 40 *µM* P301L tau containing 40 *µg/ml* polyU. Hence, we set out to quantify the rate of nucleation inside the condensates from the videos recorded using TIRF microscopy.

In the TIRF microscopy images, the nuclei (or the early-stage fibrils growing from them) appear as bright spots against the fluorescent background of the condensates (Figure 3A). Hence, automation of the detection and counting of the nucleation events within the condensates is not straightforward. To circumvent this, we adopted the following strategy. First, we create masks for all the condensates in an image by using Segment Anything Model (SAM).^36^ Then, we apply the individual masks to cut out the individual condensates. Following threshold filtering and erode-dilate processing, we could clearly distinguish the bright spots representing the nucleation events inside the condensates from the background. The process of counting the nuclei inside the individual condensates is shown in Figure 3A. To examine how the rate of nucleation depends on the size of the condensates, we computed the histograms of the number of nuclei as a function of the diameter of the condensates at various time points (see Supplementary Video 4). Then we categorized the condensates into three groups: large (diameter *>* 2.6 *µm*, magenta), medium (1.8 *<* diameter *<* 2.6 *µm*, green), and small (diameter *<* 1.8 *µm*, blue), as illustrated in Figure 3B-D. We observed that the rate of nucleation per unit area is invariant among all the three categories of the condensates (Figure 3B). In contrast, the rates of nucleation per unit volume or per condensate are not invariant with respect to the size of the condensates (Figure 3C-D). To extract a scaling law, we categorized the condensates into 12 groups by their sizes. For each group, the number of nuclei per condensate increases over time, allowing a nucleation rate to be estimated for each group (Supplementary Figure S3). The log-log plot of the nucleation rate versus condensate radius is plotted in Figure 3E. A linear fit of this data yields a scaling exponent equal to 1.9, suggesting that the rate of nucleation is proportional to the surface area of the condensates. This implies that the surface of the condensates plays a critical role in initiating the nucleation of the fibrillar aggregates. Since the imaging depth in TIRF microscopy is just about 200 nm, the nuclei counts presented above come from the basal region of the liquid droplets only. Next we performed 3D imaging of the droplets using SoRa microscopy. This approach allowed us to measure the nuclei number across an entire condensate (Supplementary Video 3). The nucleation rates obtained following the same approach described above are shown in Figure 3F-I and Supplementary Figure S4. As expected, imaging of the whole condensates yielded more nucleation events per condensate. However, the scaling exponent determined from the log-log plot of the nucleation rate and the condensate radius was still equal to 2. These results further support the conclusion that the nucleation rate predominantly depends on the surface area. This is further supported by a peculiar morphology of the fibrils observed at considerably low concentration (2.5 *µM*) of tau close to the saturation concentration where only a few small condensates are observed. These condensates were found to mature into circular fibrils that grew along the interface encircling the condensates (Figure 3J), highlighting the role of the interface in nucleation and growth of the fibrils.

**Figure 3:**
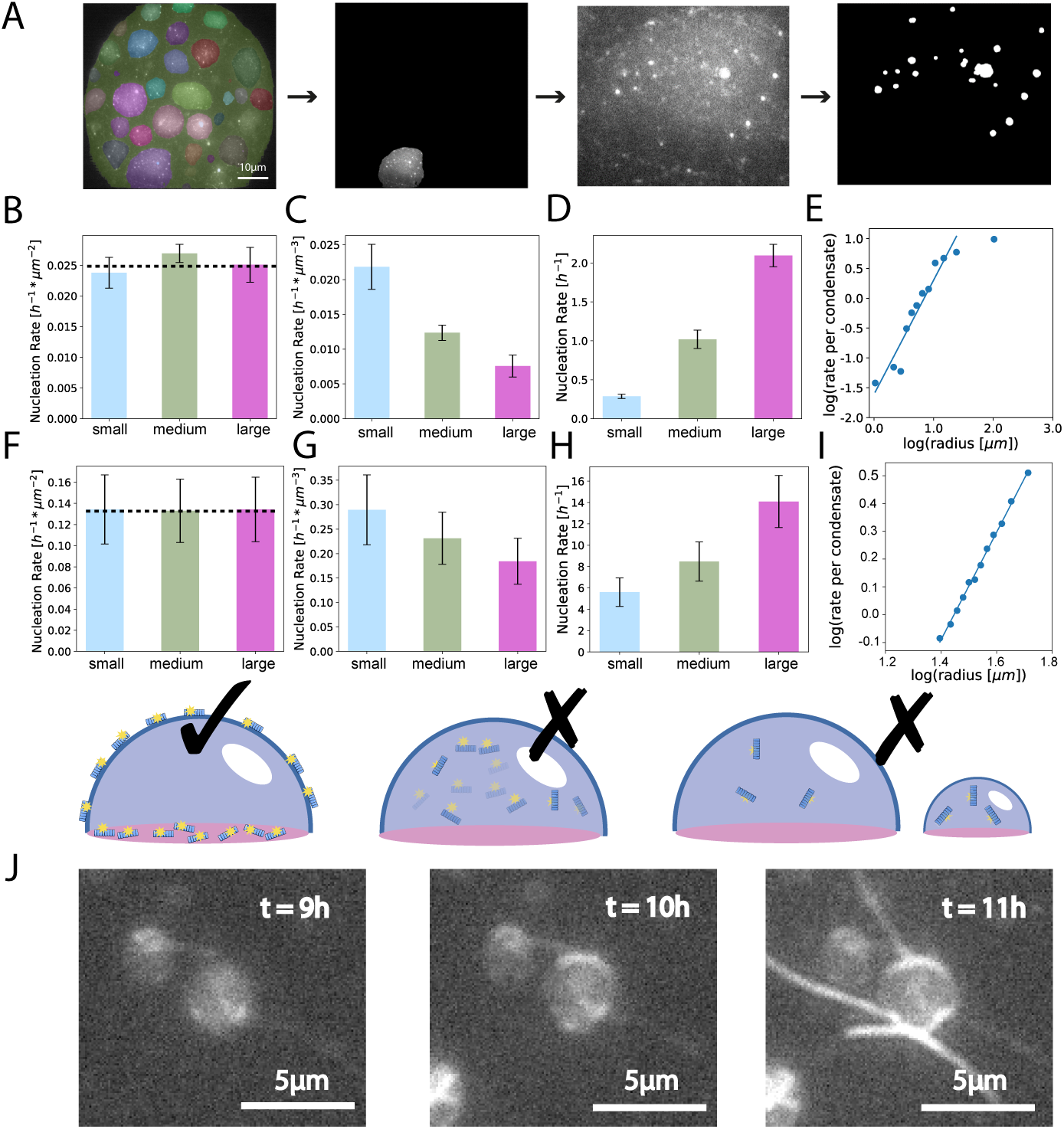
Statistics of the nucleation events in the dense phase. (A)-(E) are obtained from analysis of the TIRF microscopy recorded using 40 *µM* of P301L tau in presence of 40 *µg/ml* polyU RNA. (A) Sequence of the image processing performed to count the number of nuclei in each condensate. Each condensate in the field of view is marked with a different color in the first panel of (A). Number of nuclei per unit area of the condensates of the (B) lowest (radii 0.7 − 1.8 *µm*), (C) middle (radii 1.8 − 2.6 *µm*), and (D)largest (radii 2.6 − 37.3 *µm*) size range. (E) Nucleation rate per condensate versus condensate radius in a *log*_10_ − *log*_10_ plot. The solid line is a linear fit with slope ≈ 1.9. (F)–(I) are similar to (B)–(E) but were obtained using the SoRa experiment, which measures the total number of nuclei in the entire bulk by analyzing slices at all heights through the condensates. The slope in panel (I) is 2. The panel below (F)–(H) is a schematic representation of probable nucleation scenarios, such as at the surface or inside the condensates, and the certain number of nuclei per condensate. (J) Shows nucleation and elongation of the fibrils at the interface of small condensates at 2.5 *µM* of P301L tau in the presence of 40 *µg/ml* polyU.

### Viscosity and surface tension of the condensates during aging

To examine the liquid-to-solid transition we set out to measure the viscosity of the liquid inside the condensates and monitor its changes during the aggregation of P301L tau. Viscosity in the crowded environment within a condensate is known to be dependent on the size of the probe. ^37^ However, the viscosity of relevance here is that is experienced by the molecules of tau inside the condensates. Hence, we have used single-molecule tracking of the individual tau molecules inside the condensates to measure viscosity. To this effect, we added 50 *pM* of Alexa647 labeled tau into the solutions of the unlabeled tau mixed in the presence of polyU. TIRF microscopy images were captured at intervals of every 200 *ms* or 300 *ms* to create the videos that clearly show the movements of the single molecules of fluorescently labeled tau within the condensates (Supplementary Video 2). To analyze the videos we employed the following strategy. First, the Segment Anything Model^36^ was used to create masks for all the condensates in the first frame. Since the entire video was recorded within a 10-second time window, we assume that the position, size, or shape of the condensates did not change significantly during this duration. Hence, the mask from the first frame was applied to all the subsequent frames. These processes are illustrated in Figure 4A (materials and methods). Analysis of the trajectories shows that the mean square displacement (MSD) of the particles varies linearly with lag time on a log-log plot between 0 − 4 seconds (Figure 4B). The MSD data are then fit with the function *MSD* = *A* · *t^n^*, where 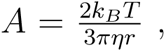 with *k_B_* being the Boltzmann constant, *T* representing temperature, *r* ≈ 1.8 *nm*^38^ denoting the average particle radius, and *η* as the viscosity of the liquid within the condensates. The value of the exponent (*n*) is found to be 0.8 ± 0.04, consistent with behavior reported previously for highly crowded milieus.^39–43^ This observation suggests that within the condensates, tau molecules exhibit crowding-induced subdiffusive motion. The viscosity of the liquid inside the droplets was estimated to be 5-6 *Pa* · *s* (Figure 4C), which is about 3-4 orders of magnitude higher than that of water, and is a value of a similar order of magnitude compared to other observations in condensates.^44^ Slow diffusivity indicates that the tau molecules experience a net attractive interaction inside the condensates.^45,46^ The slow diffusion also provides a rationalization for the slow elongation of the fibrils within the condensates, as this process depends on the mass transport of new molecules from the bulk of the condensate to the site of elongation. ^47^ Furthermore, we observed that the viscosity of the condensates is independent of the initial tau concentration, consistent with the weak dependence of the density of the dense phase with the overall protein concentration characteristic of phase separating systems. Surprisingly, however, our results reveal that the viscosity is also constant throughout the aggregation process, implying that the liquid phase remains unchanged during ageing of the condensates. This observation implies co-existence of a liquid state and a growing solid state, and suggests an aggregation pathway where solid aggregated phases start to nucleate and grow in the background of a liquid phase which is thus increasingly confined to the pores of the growing fibril gel network, but which maintains its viscosity throughout the process. Consistent with this idea, the MSD of the molecular trajectories reached a plateau at around 7 seconds (Figure 4B). The scaling exponent of the MSD curve is 0.83, indicating subdiffusive behavior, and the scaling is consistent with previous reports.^41,42,48,49^ The square root of the plateau value can be used to estimate the fibril network pore size. In these condensates this size is approximately 0.6-0.7 *µm*. This value is consistent with direct observations of the fibril network within the droplets by using Scanning Electron Microscopy (SEM) (Supplementary Figure S5). Similar inhomogeneous spatial organization at the interior of the condensates of the prion like low complexity domain (PLCD) protein was observed in the Computational studies.^50^

**Figure 4:**
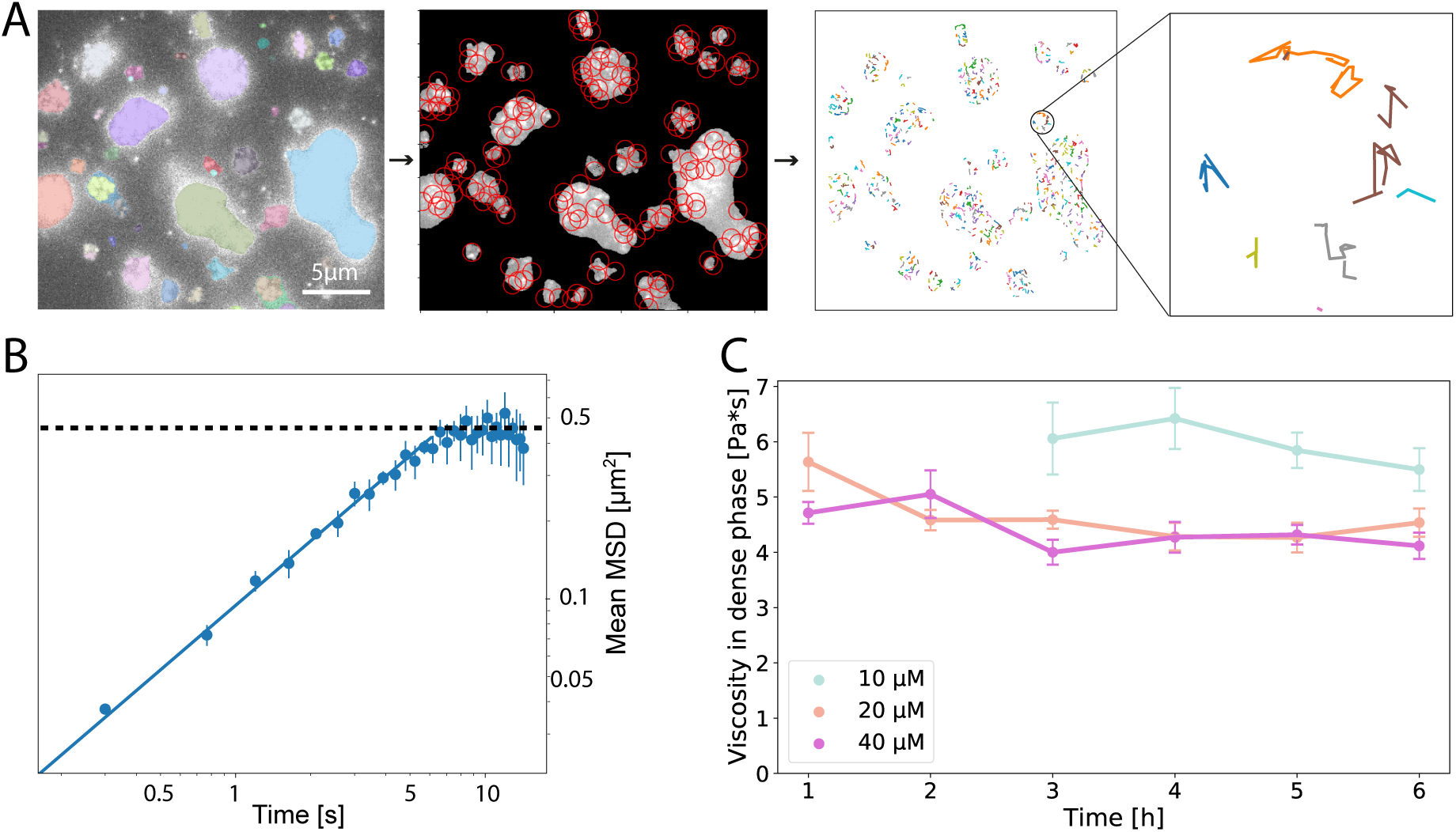
Viscosity of the dense phase measured using single-molecule tracking. (A) The sequence of image processing steps performed to analyze the data. (B) The mean square displacement (MSD) of the traced particles versus time at 4 hour. The filled circles represent MSD and the solid line is the best linear fit for 0 − 16 seconds in this *log*_10_ − *log*_10_ plot. Tau concentration is 40 *µM*. (C) The viscosity of the condensates at different time points calculated from the linear fit in B. The concentrations of P301L tau are 10 *µM*, 20 *µM* and 40 *µM*, and polyU RNA is 40 *µg/ml*.

This prompted us to explore the internal mobility properties of the aging condensates further by Fluorescence Recovery After Photobleaching (FRAP). FRAP measurements were performed using 25 *nM* fluorescein-labeled tau mixed in the presence of 10, 20, or 40 *µM* unlabeled tau and 40 *µg/ml* polyU (Figure 5A). FRAP was monitored at different stages of aging, and the half-time (*t*_1_*_/_*_2_) was estimated from the recovery kinetics (Figure 5B). Diffusion coefficient *D* of tau inside the condensate was estimated using the Soumpasis equation: ^51^ 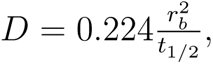 where *r_b_* is the radius of the bleached area. Then viscosity *η* of the condensates was estimated from the diffusion coefficient *D* of tau using the Stokes-Einstein equation: *D* = *k_b_T/*6*πηr*, where *r* ≈ 1.8 *nm* is the radius of P301L tau.^52^ In agreement with the observations above, we find that the viscosity of the condensates remains invariant with respect to the initial concentrations (10, 20, 40 *µM*) of tau and crucially remains constant during the ageing process (1 *h* to 24 *h*), as shown in Figure 5C. The viscosity values obtained from the FRAP measurements (*η* = 1.5 to 3.5 *Pa* · *s*) agree well with those obtained using the single-particle tracking presented in Figure 4C. Crucially, the FRAP measurements further revealed that the fraction of the recovered fluorescence decreased considerably with maturation. Therefore, while the viscosity of the droplets remained unchanged, the fraction of the immobile fractions increased with time, consistent with the picture in which a solid gel network grows within the initially liquid condensate as a function of time.

**Figure 5:**
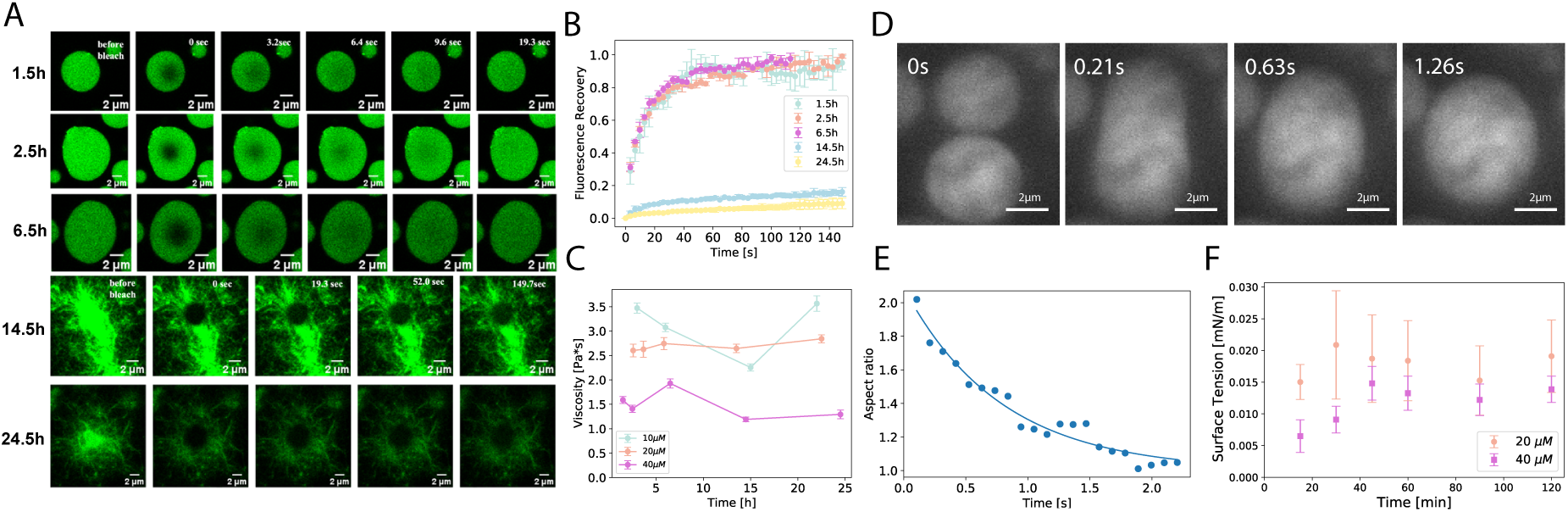
Viscosity and surface tension measured using FRAP and confocal microscopy. (A) Confocal microscopy images of the condensates during the FRAP experiment. (B) FRAP recovery kinetics at different stages of ageing of the condensates. (C) The viscosity of the condensates at different concentrations of P301L tau and maturation time. (D) Fusion of two condensates captured by epifluorescence microscopy. (E) Representative aspect ratios of condensates over time are shown for a monomer concentration of 20*µM*, measured at 15 minutes. (F) Surface tension of condensates at different time points and different monomer concentrations.

Since we observed that the surface of the condensates plays a key role in fibril nucleation, we set out to measure their surface tension using epifluorescence microscopy. To do so, we focussed on the fusion dynamics of the droplets, which reflect the relationship between the condensate viscosity and surface tension. Figure 5D presents representative sequential epifluorescence microscopy images illustrating the fusion process of two condensates in liquid. The time course of the change in the aspect ratios of the condensates during fusion was fitted by an exponential curve (Figure 5E) to determine the fusion time, *τ_fusion_*.^53,54^ The ratio of droplet viscosity to surface tension, known as the inverse capillary velocity, was estimated using the relationship *τ_fusion_* = (*η/γ*) · *l*, where *η* is the viscosity, *γ* is the surface tension and *l* is the characteristic length of the condensate. The viscosity of the condensate was measured using particle tracking (Figure 4C) and FRAP (Figure 5C). The estimated surface tension from these measurements is about 0.01 − 0.02*mN/m*, similar to the values reported earlier for condensates of other proteins.^44^ The calculated surface tension of the condensates at different time points and monomer concentrations (20*µM* and 40*µM*) is shown in Figure 5F. We observed that the surface tension of the condensates remained unaffected by tau concentration and remained constant over time (t ¡ 2 hours). Surface tension was not measured at times beyond 2 hours (t ¿ 2 hours) due to the rarity of the fusion events.

## Conclusion

The liquid-to-solid transition of the biomolecular condensates plays crucial roles in neurodegenerative diseases, however, molecular determinants of this process are still poorly understood. We have measured the rates of the key processes that drive this transition. We showed that TIRF microscopy is a powerful technique to visualize the amyloid nuclei and the fibrils on the liquid condensates of tau. We were able to measure the rate of nucleation on the individual condensates and observed that it is proportional to *r*^2^, where *r* is the radius of the condensates. Hence, our results indicate that the surface of the tau condensates must be the sites for at least the initial nucleation step. This differs from what is expected in the case of homogeneous nucleation where the rate of nucleation is proportional to the volume (∼ *r*^3^) of the droplets.^55^ Although the high concentration of tau inside the liquid droplets is expected to promote homogeneous primary nucleation in bulk, we think that the high surface-to-volume ratio of the condensates dominates in promoting primary nucleation on the condensate surface.^56^ Computational studies have shown that the conformation of the protein molecules at the interface exhibit certain unique features, e.g, the molecules are extended and aligned perpendicular to the interface.^50^ Strong alignment of the protein monomers at the interface could be responsible for promoting nucleation. Furthermore, we were able to trace the elongation of the individual fibrils and determine their rate of elongation. Our measurements show that the rate of elongation is approximately 4 *µm/h* both inside the condensate and in the dilute phase although the concentration *ϕ* of tau is about 70 times higher in the dense phase than in the dilute phase. However, the effective diffusion of the tau molecules is about 5000 -times slower inside the condensates than in the dilute phase. Therefore, the rate *k* of elongation of the fibrils inside the condensates is about (5000/70 =) 71 times faster than expected. The rate of a reaction in a crowded environment may be reduced due to slow diffusion but it can also be enhanced due to excluded volume and confinement effects.^34,57,58^ Therefore, a complex interplay of the intermolecular interactions (tau-tau, tau-RNA and RNA-RNA), and crowding effects govern the growth of the fibrils within the dense milieu of the condensates. Finally, single particle tracking and FRAP measurements revealed that ageing of the condensates is characterized by coexistence of a growing solid phase within the liquid phase. The growth of the solid phase is mediated by nucleation and the elongation of the amyloid fibrils. The viscosity of the liquid inside the condensates remains nearly unchanged even during aging, although the fraction of the mobile phase decreases. Therefore, at least in the case of P301L tau the liquid to solid transition does not alter the internal friction and intermolecular forces of the liquid inside the condensate. Finally, we propose that the methodologies used here will help in the characterization of the liquid-to-solid transitions of other proteins involved in the neurodegenerative diseases.

## Materials and methods

### Protein expression and purification

In all experiments, we used the truncated variant of 2N4R tau (residues 255–441), which contains the disease relevant mutation P301L along with two additional mutations (C291S/C322S) to remove the native cysteine residues and a 6x His Tag tag at the N-terminal to facilitate purification of the protein. The plasmid was a kind gift from Songi Han lab (University of California, Santa Barbara). The detailed protocol for expression and purification of the protein can be found elsewhere.^17^ Briefly, the protein was expressed in E. coli BL21(DE3) cells, and purified using Ni-NTA column using standard Ni-NTA purification protocol. Finally, the purified protein was subjected to buffer exchange through dialysis in 20 mM HEPES buffer, pH 7.4, utilizing a 10 kDa dialysis bag. Purity was validated via SDS-PAGE and mass spectrometry analysis. The concentration was determined by UV-Vis absorbance at 277 *nm* employing a molar extinction coefficient of 2980 *cm^−^*^1^*M^−^*^1^. For labeling tau with a fluorophore, P301L tau containing a single cysteine residue (viz, C291S/P301L/C322) was used. Labeling was performed using either Alexa647 maleimide or Alexa488 maleimide. The labeled protein was purified by size exclusion chromatography by using a Superdex 200 column (GE Healthcare).

### Sample Preparation for the Microscopy Experiments

A multiwell chamber (4-well or 96-well coverglass-bottom chamber, Celvis) was cleaned with 10M NaOH and thoroughly rinsed with MilliQ water to eliminate dirt from the glass surface, to ensure high-contrast TIRFM imaging.^31^ The wells were disinfected using a 70% (v/v) ethanol-water solution and subsequently rinsed with MilliQ water. Concentrated P301L tau stock was diluted to the desired concentrations into 20 *mM* Hepes buffer at pH 7.4, containing 10 *µM* ThT, 10 *mM βMe*. polyU (40 *µg/ml*) RNA was added to induce liquid-liquid phase separation prior to recording the images in a home-built TIRF microscope. Details of the microscopy setup can be found elsewhere.^31^ Formation of the liquid condensates and the amyloid fibrils were monitored using ThT fluorescence. We recorded the nucleation events within condensates in three dimensions using a *Nikon Ti2 Eclipse CSU-W1 SoRa* (Japan) spinning disk super resolution by optical pixel reassignment (SoRa) microscope, with samples prepared in the same manner as for TIRF microscopy.

The single molecule tracking experiment was performed using Alexa647 labeled P301L tau. 100pM Alexa647 labeled tau was added to the solution of unlabeled P301L tau to record the videos of the single molecules in the TIRF microscope. The images were captured with a fast SCMOS camera (PCO). A confocal microscope (Olympus) was used for recording Z-stack images of the droplets and for performing the FRAP (Fluorescence Recovery After Photobleaching) measurements. In these experiments the samples were supplemented with 25 nM Alexa-488 labeled P301L tau. FRAP kinetics were utilized to measure viscosity of the droplets.

### Sample Preparation for SEM Analysis

The sample preparation protocol closely resembles that used for TIRFM experiments, the main distinction being the inclusion of a carefully cleaned p-type silicon wafer as the substrate. A small piece of silicon wafer was placed on the glass surface within a single-well chamber. The tau and polyU mixture was then added, ensuring uniform coverage of the silicon wafer surface. The assembly was incubated at 25*^◦^*C for 24 hours to promote fibril formation directly on the wafer substrate. Following incubation, the silicon wafer was gently immersed in a Milli-Q water bath for 10 seconds to remove residual buffer and unbound material. The wafer was subsequently air-dried under ambient conditions. The dried silicon wafer was then subjected to scanning electron microscopy (SEM) to resolve the fibrillar architecture at nanometer-scale resolution.

### Image Analysis

To trace elongation of the fibrils, the TIRF microscopy image sequences were imported into Fiji/ImageJ.^59^ Every frame in this image sequence was then accessed using NeuronJ,^60^ an ImageJ plugin initially created for tracing neurites. Within this plugin, we manually identified and selected the fibrils by employing the inbuilt Livewire segmentation algorithm. The length of each traced fibril was recorded and plotted in Figure 2. When masking the condensates in Figure 3 and Figure 4, we utilized AI tools, specifically the “segment anything” tool developed by Meta.^36^ We employed the default Sam’s model to segment all condensates in every frame. We also use the same protocol to track the nucleation events in SoRa images. The volume of each condensate is the sum of the area of each slice of the condensate multiplied by the distance between two slices, which is 0.6*µm*. 2849 condensates have been measured and analyzed for TIRF experiments, and 1782 condensates have been measured and analyzed for SoRa experiments. The Python package ‘trackpy’ was employed^61^ to automatically track all the bright particles and record their trajectories, as shown in Figure 4A.

## Acknowledgement

The authors thank Dr. Yann Fichou for sharing the plasmid of tau protein with us. This work received generous support from the Department of Atomic Energy, Government of India (project ID no. RTI 4007 to M.S.R.S. and K.G.), the Science and Engineering Research Board, Government of India (grant no. CRG/2020/005527 to K.G.), the ERC grant DiProPhys (agreement ID 10100161, T.P.J.K), and the Nidus studentship scheme (J.W.).

## List of Abbreviations

**A***β*: Amyloid *β*
AD: Alzheimer’s disease
IDPs: Intrinsically Disordered Proteins
FRAP: Fluorescence Recovery After Photobleaching
LLPS: Liquid-Liquid Phase Separation
LST: Liquid to Solid Transition
ThT: Thioflavin T
TIRFM: Total Internal Reflection Fluorescence Microscopy
SoRa: Spinning Disk Super Resolution by Optical Pixel Reassignment
SAM: Segment Anything Model

