## Supplementary text for "Liquid-solid coexistence at single fibril resolution during tau condensate ageing"

### List of Contents

**Section S1**: Elongation rates at different concentrations of tau and Supplementary figure S1.

**Section S2**: Comparing concentration of tau inside and outside of the dense phase and Supplementary Figure S2.

**Section S3**: Rate of nucleation in the dense phase and Supplementary Figure S3 and S4.

**Section S4**: SEM of condensates.

**Section S5**: Growth of fibrils in the dilute and the dense phases.

**Supplementary video 1**: The stop-and-go aggregation kinetics of P301L tau captured using TIRF microscopy.

**Supplementary video 2**: Single particle movement recorded with a frame interval of 300ms inside P301L tau condensates using TIRF microscopy.

**Supplementary video 3**: SoRa microscopy images of P301L tau condensates taken at 90 minutes.

**Supplementary video 4**: Number of nuclei at different size ranges of the condensates. The nuclei number distribution at different time points.

**Supplementary Figure S1**: Fibril elongation traces in the dilute phase and the dense phase.

**Supplementary Figure S2**: Comparing tau concentration inside and outside of the dense phase.

**Supplementary Figure S3**: Nucleus number per condensate versus time for different sizes of condensates measured by TIRF.

**Supplementary Figure S4**: Nucleus number per condensate versus time for different sizes of condensates measured by SoRa.

**Supplementary Figure S5**: Analyzing SEM images to estimate the pore size of the condensates.

### Supplementary Information

#### S1. Elongation rates at different concentrations of tau

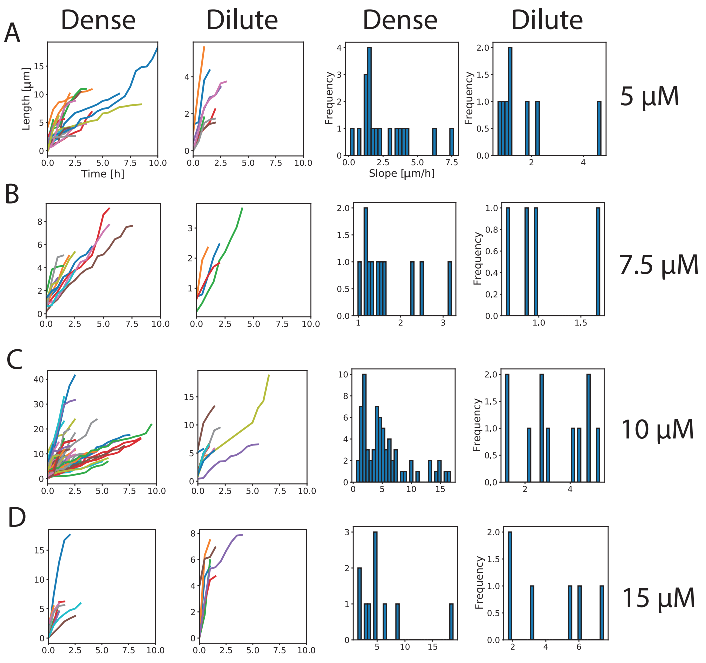


**Supplementary Figure S1: Fibril elongation traces in the dilute phase and the dense phase.** (A)-(D) show the time traces of the individual fibrils and distributions of the elongation rates of traced fibrils in the dilute and the dense phase. The concentration of P301L tau varies from $5 - 15$ $\mu M$ in the presence of 40 $\mu g/ml$ polyU. The paused sections of the stop-go elongation behavior have been removed. The elongation rate is the slope of the linear fit to the remaining fibril length over time data.

#### S2. Comparing concentration of tau inside and outside of the dense phase

To quantify the concentration of tau, we mixed 50 nM Alexa488-tau with 10 $\mu$M unlabeled tau and 40 $\mu$g/ml polyU, and incubated the sample at RT for 6 hr. Then we measured the fluorescence intensity of Alexa488 tau inside the dense and the dilute phases using a fluorescence correlation spectroscopy (FCS) setup. We observed that prolonged exposure to light causes photobleaching of the Alexa488-tau inside the dense phase. To minimize photobleaching we used a motorized XY sample stage and recorded the fluorescence intensity from the sample while the stage was moving continuously. Since FCS uses a confocal detection scheme, fluorescence intensity was being recorded from inside and outside of the dense phase/liquid droplets of tau as the sample stage was scanning different regions of the sample. Hence, the fluorescence intensity was found to vary dramatically (See Supplementary Figure S2). The Low and the high values of the intensities came from the dilute and the dense phases respectively. In contrast, fluorescence intensity measured from 50 nM Alexa488-tau in absence of polyU, i.e., in absence of phase separation remained constant during scanning of the sample stage (data not shown). The ratio of the concentration of tau inside and outside of the droplets $((C_{dense}/C_{dilute}))$ is estimated from the ratio of the fluorescence of Alexa488-tau inside and outside of the droplets $(F_{dense}/F_{dilute})$, i.e., $(C_{dense}/C_{dilute}) = (F_{dense}/F_{dilute})$. Our results suggest that the dense phase is about X/Y = 70 times more concentrated than the dilute phase.

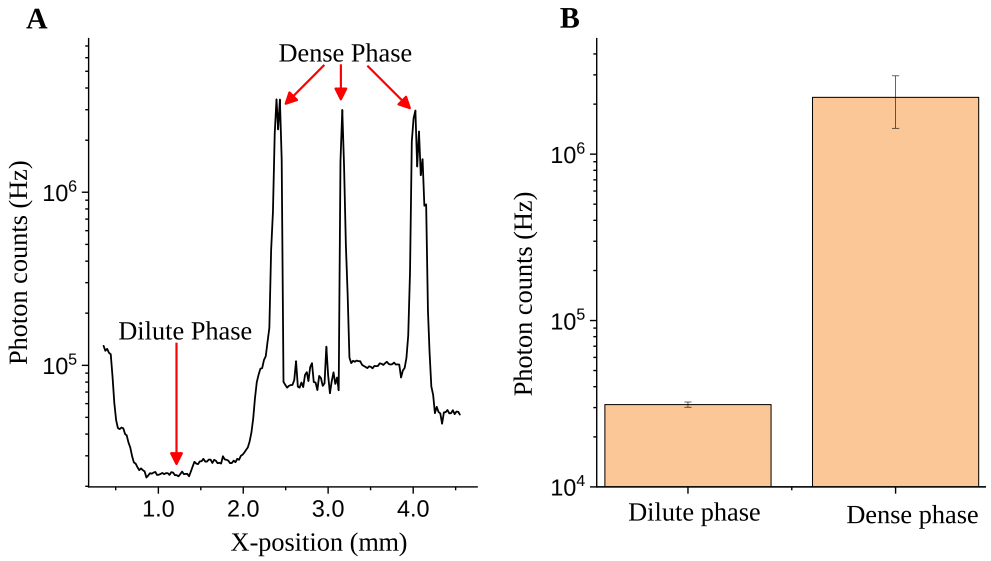


**Supplementary Figure S2: Comparing tau concentration inside and outside of the dense phase.** (A) Fluorescence was recorded from the sample containing 50 nM Alexa488-tau, 10 $\mu$M unlabeled tau and 40 $\mu$g/mL polyU while the sample stage was scanning continuously in the X-direction. (B) Average fluorescence intensities of the dilute and the dense phases were determined from the regions of the minima and the maxima of the intensity trace shown in A. The ratio of dense phase to dilute phase concentration is measured to be $(C_{dense}/C_{dilute}) = (F_{dense}/F_{dilute}) \approx 70$.

#### S3. Rate of nucleation in the dense phase

For 40 $\mu g/ml$ of P301L tau with 40 $\mu g/ml$ polyU, at different time points, we counted the number of nuclei at different size ranges of the condensates. We plot the nuclei number distribution at different time points, as shown in Supplementary video 4.

Nuclei number per condensate versus time for 12 groups of condensates are shown in Supplementary Figure S3 (TIRF) and Supplementary Figure S4 (SoRa).

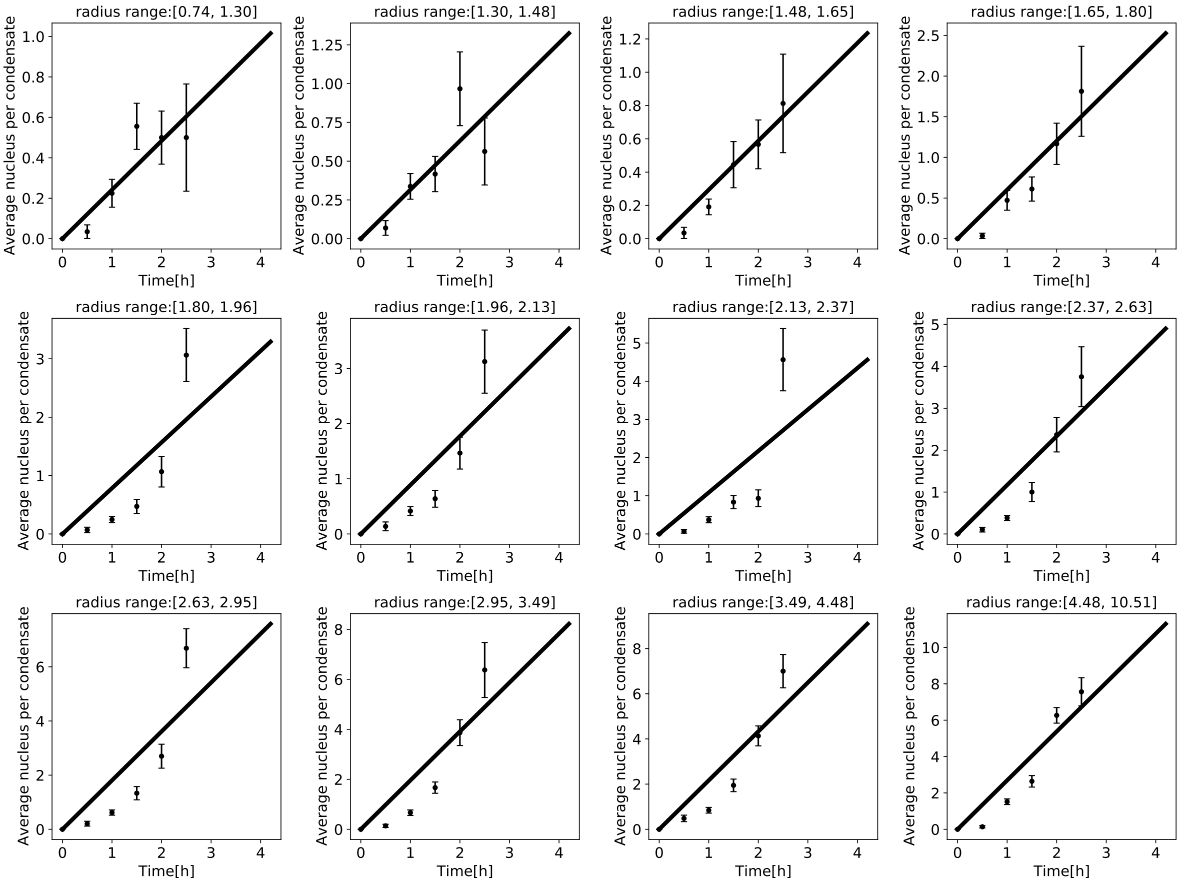


**Supplementary Figure S3: Number of nuclei per condensate versus time for condensates of different sizes measured from tirf.** The condensates are separated into 12 groups according to their size. A linear fit has been applied for each group of condensates in their average nucleus per condensate versus time plot. It may be seen that the slope of the line increases from 0.25 condensate/h to 2.5 condensate/h with the increase of radius from 1.3 to 4.5 $\mu$m.

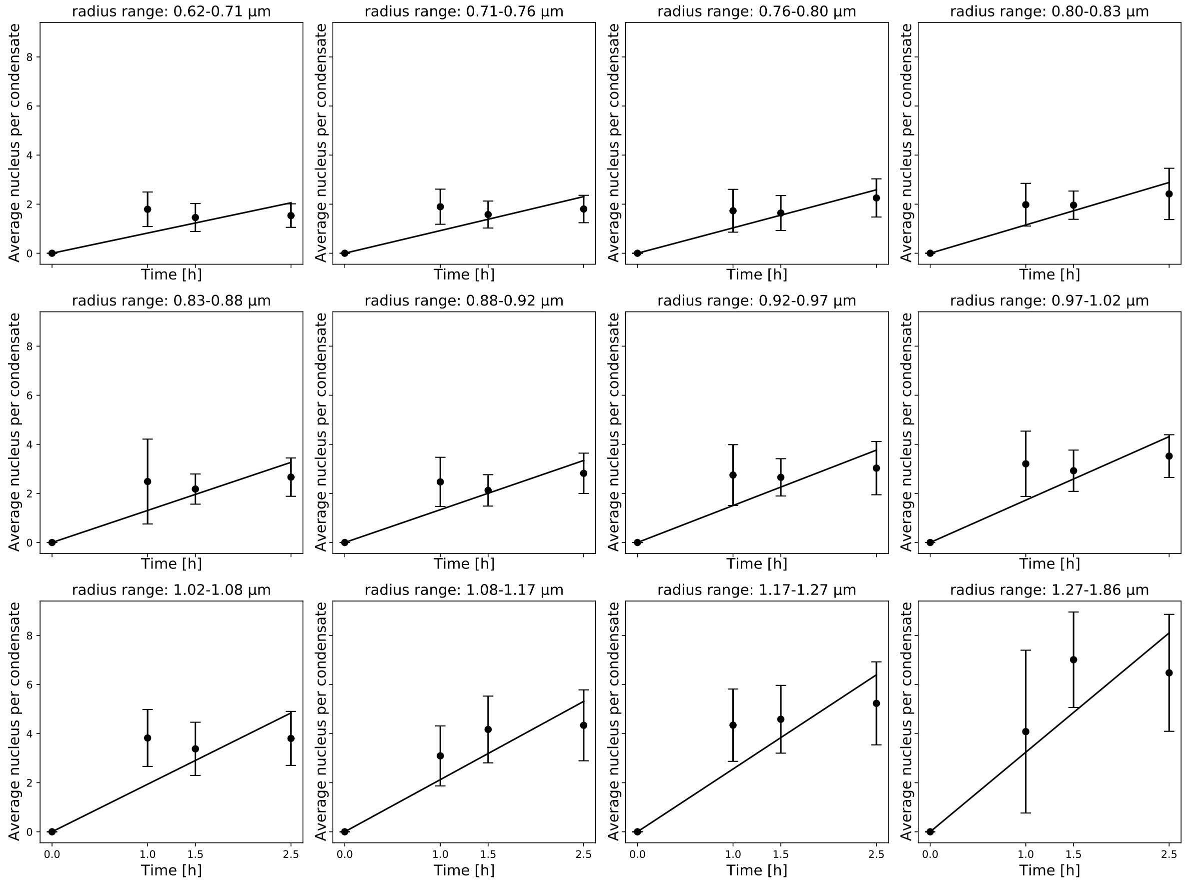


**Supplementary Figure S 4: Number of nuclei per condensate versus time for condensates of different sizes measured from SoRa.** The condensates are separated into 12 groups according to their size. A linear fit has been applied for each group of condensates in their average nucleus per condensate versus time plot.

#### S4. SEM of condensates

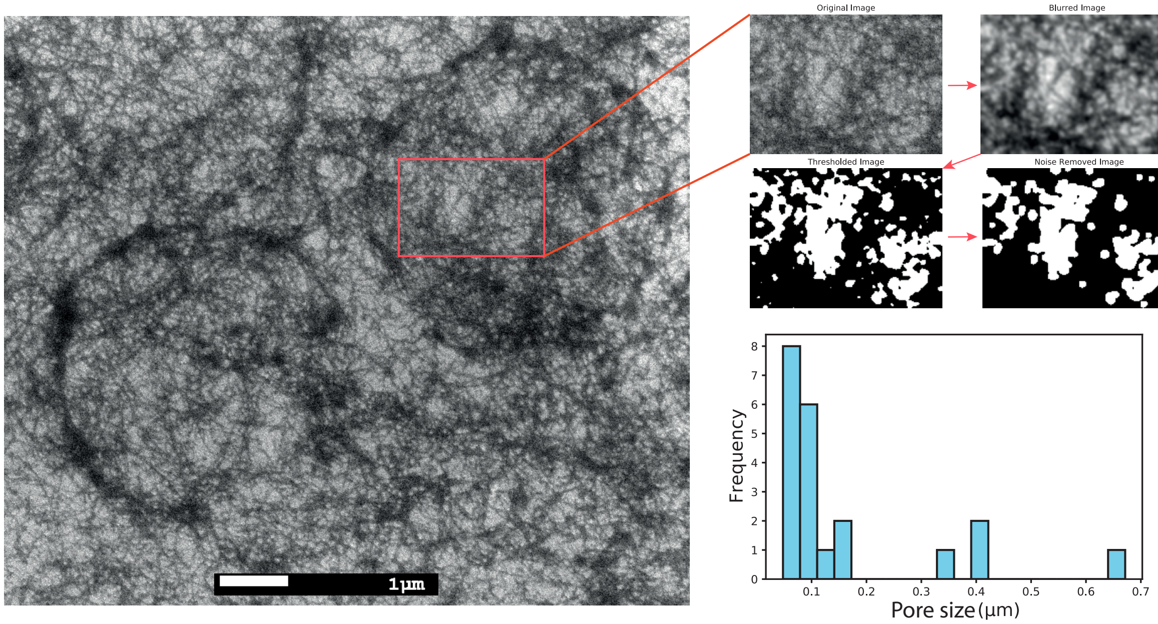


**Supplementary Figure S5: Using SEM image to measure the distribution of condensates pore sizes.**

#### S5. Growth of fibrils in the dilute and the dense phases

Supplementary Video 1 illustrates the stop-and-go aggregation kinetics of tau protein in a sample containing 5 $\mu M$ P301L tau and 40 $\mu g/ml$ polyU. The arrows in the video indicates the fibril that shows a pause and then starts its elongation.
